# The ability of single genes vs full genomes to resolve time and space in outbreak analysis

**DOI:** 10.1101/582957

**Authors:** Gytis Dudas, Trevor Bedford

## Abstract

Inexpensive pathogen genome sequencing has had a transformative effect on the field of phylodynamics, where ever increasing volumes of data have promised real-time insight into outbreaks of infectious disease. As well as the sheer volume of pathogen isolates being sequenced, the sequencing of whole pathogen genomes, rather than select loci, has allowed phylogenetic analyses to be carried out at finer time scales, often approaching serial intervals for infections caused by rapidly evolving RNA viruses. Despite its utility, whole genome sequencing of pathogens has not been adopted universally and targeted sequencing of loci is common in some pathogen-specific fields. In this study we aim to highlight the utility of sequencing whole genomes of pathogens by re-analysing a well-characterised collection of Ebola virus sequences in the form of complete viral genomes (~19kb long) or the rapidly evolving glycoprotein (GP, ~2kb long) gene. We quantify changes in phylogenetic, temporal, and spatial inference resolution as a result of this reduction in data and compare these to theoretical expectations. We propose a simple intuitive metric for quantifying temporal resolution, *i.e.* the time scale over which sequence data might be informative of various processes as a quick back-of-the-envelope calculation of statistical power available to molecular clock analyses.

## Introduction

The combination of decreasing cost of sequencing and the unparalleled insight it offers have led to the adoption of pathogen genetic sequencing as one of the most effective tools in a modern epidemiologist’s toolkit. When coupled with sophisticated models of evolution, pathogen sequences can be used to look into epidemiological features such as cryptic transmission (Faria et al., 2017), migration (Lemey et al., 2009, 2014), and origins (Smith et al., 2009) of infectious diseases, amongst others. Pathogen sequences also contain information about past temporal dynamics before sequence data have been collected (Raghwani et al., 2012) due to the pattern of shared and unique mutations inherited from preceding generations. Molecular phylogenetic approaches rely on decoding these patterns of shared mutations into a nested graph known as the phylogenetic tree. Pathogens often have short generation times and some, like RNA viruses, also possess polymerases with low replication fidelity such that mutations are generated at a rapid pace (Drummond et al., 2003; Biek et al., 2015) leading to fast differentiation of pathogen lineages at the genetic level as they spread. With appropriate sampling and information (“metadata”) about sequences, historic population dynamics can be inferred and quantified from pathogen phylogenies. Changes in pathogen population sizes over time (Pybus et al., 2000), inference of unobserved ancestral states (Lemey et al., 2009; Dudas et al., 2018), correlates of processes (Faria et al., 2013; Lemey et al., 2014; Dudas et al., 2017), and overall phylodynamic (Grenfell et al., 2004) patterns can be inferred from molecular phylogenies and used to understand patterns of pathogen transmission at a number of scales.

Before widespread adoption of high-throughput sequencing limitations and costs led to amplification and sequencing of short fragments of pathogen genomes (Jin et al., 1999, 2005). These subgenomic fragments were often chosen for their diversity, such as viral surface glycoproteins that experience selective pressures from vertebrate immune systems, or their utility, such as routine sequencing of HIV *pol* gene to test for drug resistance (Kaye et al., 2008; Rhee et al., 2003). Whilst subgenomic fragments of pathogens are very accurate and specific as diagnostic markers and informative about long-term evolution, their length (dictated by the compromise between information content and ease of sequencing) limits their utility in detailed molecular epidemiology investigations, for example during outbreaks (Wohl et al., 2018), as only mutations occurring within the small region of the genome that is sequenced are available for phylogenetic inference.

Molecular clocks have been particularly useful in molecular epidemiology, where the accumulation of mutations between sequences is used as a noisy approximation for elapsed time, given either times of events in the phylogeny (sequence dates or dates of common ancestors) or a previously determined molecular clock rate. Generally, neutral pathogen variation at the nucleotide level ebbs and flows under the forces of population genetics, unlike beneficial or deleterious variation which tends to either fix or be purged rapidly, respectively. Due to their random and discrete nature, mutations are modelled as a Poisson process (Yang, 2006) where the waiting time *t* for observing a mutation at a single site is exponentially distributed with evolutionary rate parameter *R*. The probability of observing 0 mutations at a single site after time *t* is *e*^−*Rt*^ and the probability of at least one mutation is therefore 1 − *e*^−*Rt*^. Higher evolutionary rates *R* or waiting times *t* result in higher probabilities of observing at least one mutation at the site in question. Since sites are assumed to evolve independently, the probability of observing at least one mutation across *L* sites is

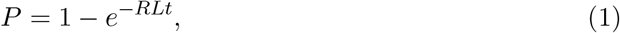

where *RL* is expressed in substitutions per year (rate in substitutions per site per year multiplied by number of sites). Under a given evolutionary rate *R*, and sequence length *L*, we can rearrange the equation to quantify the mean waiting time for at least one mutation:

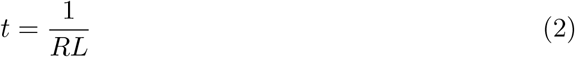

When the evolutionary rate *R* or sequence length *L* are low mean waiting times are lengthened and *vice versa*. Since both maximum plausible evolutionary rates and genome length are largely dictated by deleterious mutation load (Gago et al., 2009), neither quantity will vary substantially for a given pathogen, though individually *R* and *L* can vary substantially, where for example viruses have high *R* and low *L* on average, and bacteria have higher *L* but lower *R*.

In this study we quantify how much information relevant to phylodynamic analysis is lost when shorter genomic regions are used instead of full genomes. By focusing our attention on a subset (600 sequences) of a well-characterised genomic sequence data (comprised of >1600 viral genomes) set derived from the West African Ebola virus epidemic of 2013-2015 (Dudas et al., 2017) we estimate loss in precision and accuracy of molecular clock models and phylogeographic inference methods when only the glycoprotein gene (GP), a region representing just 10% of the viral genome, is analysed despite GP evolving at rates faster than the genomic average. Our methods rely on masking tip dates and locations for 60 (10%) of the sequences in a classic training-testing split, where we re-infer these parameters as latent variables in MCMC. We show that this reduction in data not only leads to severe convergence issues in Markov chain Monte Carlo (MCMC) analyses by removing the constraints additional data impose on plausible parameter space but can also result in unreliable tip date and location inference. Despite achieving much better temporal resolution when using complete viral genomes we still find residual error caused by inherent randomness of mutations which is close to theoretical expectations (Eq 2). We refer to this as the temporal horizon, *i.e.* a temporal resolution limit where population processes occurring at a rate faster than the rate at which mutations enter and are observed in a population will not be captured with high fidelity, even with genome sequences.

## Results

### Loss of phylogenetic signal

Figure 1 shows the reconstructed phylogenies in substitution space (right) and time space (left) for 600 complete Ebola virus genomes (top) or just GP sequences (bottom). Although higher levels of divergence are observed in the GP dataset (note that x axes between genome and GP phylogenies are not consistent), as seen from tree height, the differences in the number of non-polytomic nodes between genomic and GP data are clear, indicating substantially better resolution in disentangling the exact relationships between lineages in the former. Figure S1 shows where in the better resolved maximum likelihood phylogeny of genome sequences mutations that occurred in just the region spanning the GP gene. Internal branches of a phylogeny correspond to hypotheses of common ancestry and in the case of GP only 42 internal nodes are identified in the maximum likelihood phylogeny compared to 210 internal nodes for complete genomes. The few aspects of the West African epidemic that can be inferred from both GP and genome phylogenies, only the virus’ origins in Guinea are clear, with details of its onwards spread largely lost in the GP phylogeny. Genomic data, on the other hand, despite a reduction from over 1600 sequences described in the original study (Dudas et al., 2017) down to just 600, still contain information about the role of Sierra Leone’s epidemic in maintaining transmission across the region through both endemic proliferation of lineages and their spread to neighbouring countries.

**Figure 1.**
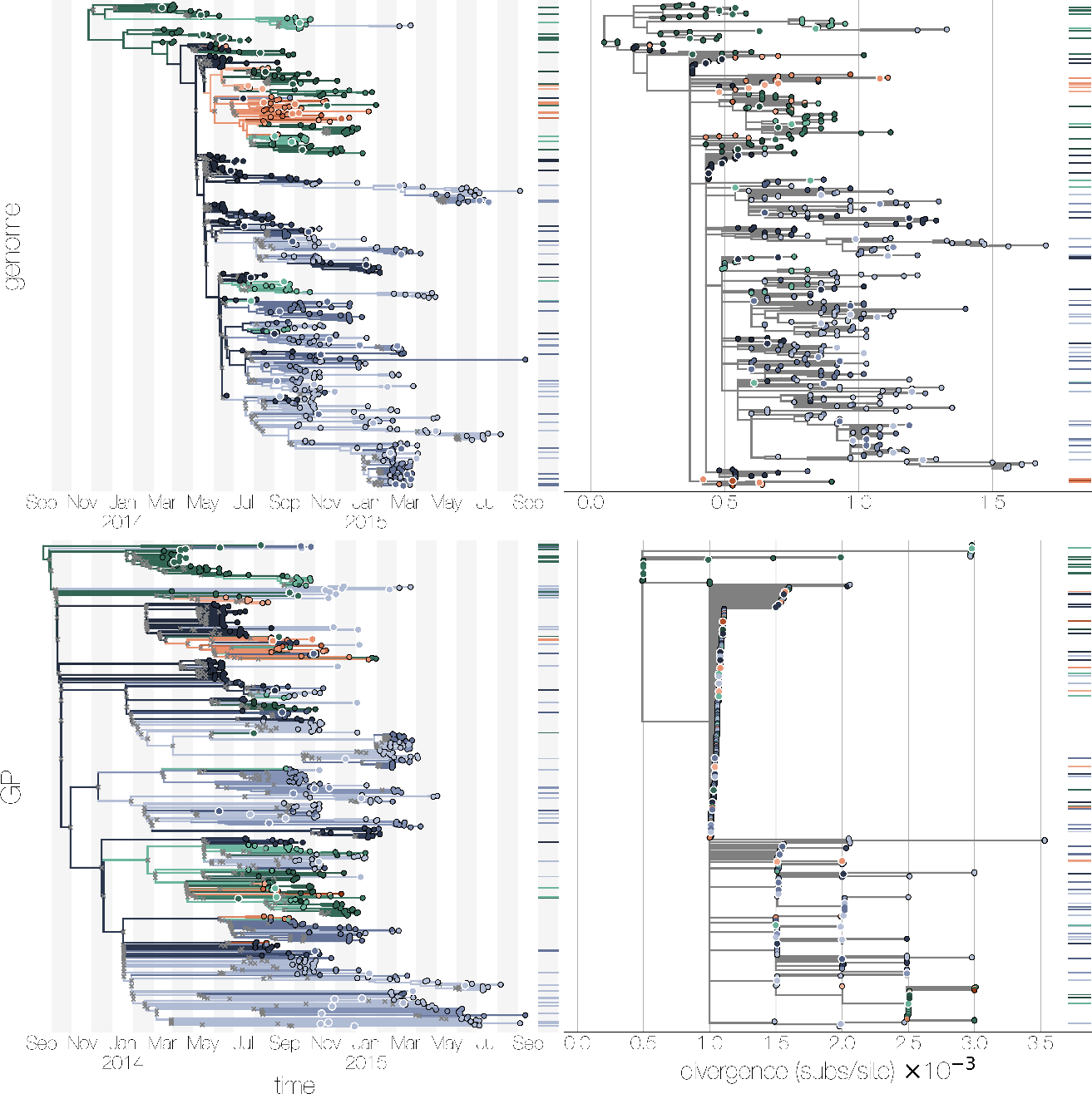
Phylogenies of West African Ebola virus genomes (top) or GP sequences (bottom). Temporal phylogenies recovered using BEAST are shown on the left, maximum likelihood phylogenies recovered with RAxML are on the right. Tips are coloured based on country (Sierra Leone in blue, Liberia in red, Guinea in green) and location (lighter colours indicate administrative divisions lying towards west of the country). Tips outlined in white indicate the 60 chosen for date and location masking, ticks to the right of phylogenies indicate the y positions of masked tips, coloured by their true location. In temporal phylogenies branches are also coloured based on GLM-inferred ancestral locations. Nodes in temporal phylogenies with <0.10 posterior probability are indicated with grey X marks. Maximum likelihood phylogenies on the right are rooted via temporal regression in TreeTime. Note that x axes between maximum likelihood phylogenies are not consistent, as GP evolves faster than the genomic average and thus has more substitutions per site.

Unlike maximum likelihood phylogenies where branch lengths are directly proportional to the expected number of substitutions, branch lengths in temporal phylogenies are usually smoothed out by the fact that a range of dates are compatible with a given number of mutations on a branch. Thus even large polytomies can be resolved into a branching structure derived from the tree prior, albeit without much support for any given configuration. This can be seen in Figure 1, where temporal phylogenies (left) appear to have similar degrees of resolution, yet the GP dataset (bottom) contains more nodes with less than 0.10 posterior support (marked by grey crosses). Similarly, there is a noticeable degree of branch clustering by country in the GP temporal phylogeny, possibly caused by proximity of locations within country, which in the absence of genetic information cannot be resolved to the same degree as with genomic data.

In contrast to the maximum likelihood phylogeny of GP on the right (Figure 1) its corresponding temporal phylogeny on the left exhibits a reconstruction of the West African epidemic largely consistent with what has been established previously (Dudas et al., 2017). This is likely to be caused by the combined effects of two sources of information. First, additional information is added by specifying the collection dates for sequences, which might exclude certain topologies from being considered during MCMC on account of the relatively small effective population size of Ebola virus in West Africa. Second, the generalised linear model approach to inferring migration is information-rich as it provides over 3000 possible parameter values (pairwise migration rates between locations) per predictor matrix, and thus if a few branches are strongly selecting for a “correct” predictor matrix to be included in the migration model that predictor matrix can then be used to determine the likely locations of branches for which less information is available. However, a simpler continuous time Markov chain model where each individual pairwise migration rate is inferred individually in a maximum likelihood framework exhibits broadly similar patterns too (Figure S2).

### Loss of temporal information

Inferring masked tip dates from 10% of the sequences (Figure 2) is an intuitive way to show both the inherent noisiness of molecular clock estimates, as reflected in the width of 95% highest posterior density intervals for inferred dates, and the differences in temporal resolution between GP and genome alignments. True collection dates for genomes are mostly (56 out of 60, corresponding to a coverage probability of 0.93) within the 95% highest posterior density of estimated dates, and the mean absolute error is ≈22 days across all masked tips. In contrast, the 95% HPDs for inferred dates in the GP dataset capture more of the true dates (58 out of 60, coverage probability ≈0.96) at the cost of markedly reduced precision, with mean absolute error going up to 106 days or 3.5 months. Despite having lower coverage probability, more precise date estimates are derived from complete genomes with an average 95% HPD width of ≈102 days, compared to ≈458 days for GP. Another way of thinking about where the loss of information occurs is to consider root-to-tip against tip date regressions shown in Figure S3, where waiting times for mutations are too long to estimate the slope of the regression reliably, as every new mutation is seen across sequences collected over a longer interval of time. Observed errors (Figure 2, but also Figure S4 for maximum likelihood equivalent) for both datasets are very close to theoretical expectations calculated using Equation 2: 22 (observed) versus 20 (expected) days for Ebola virus genomes, and 106 (observed) versus 113 (expected) days for GP. Also note that for many tips in the GP data set independent Markov chains converged onto different dates for masked tips (*i.e.* local maxima) resulting in multi-peaked posterior samples after combining independent analyses.

**Figure 2.**
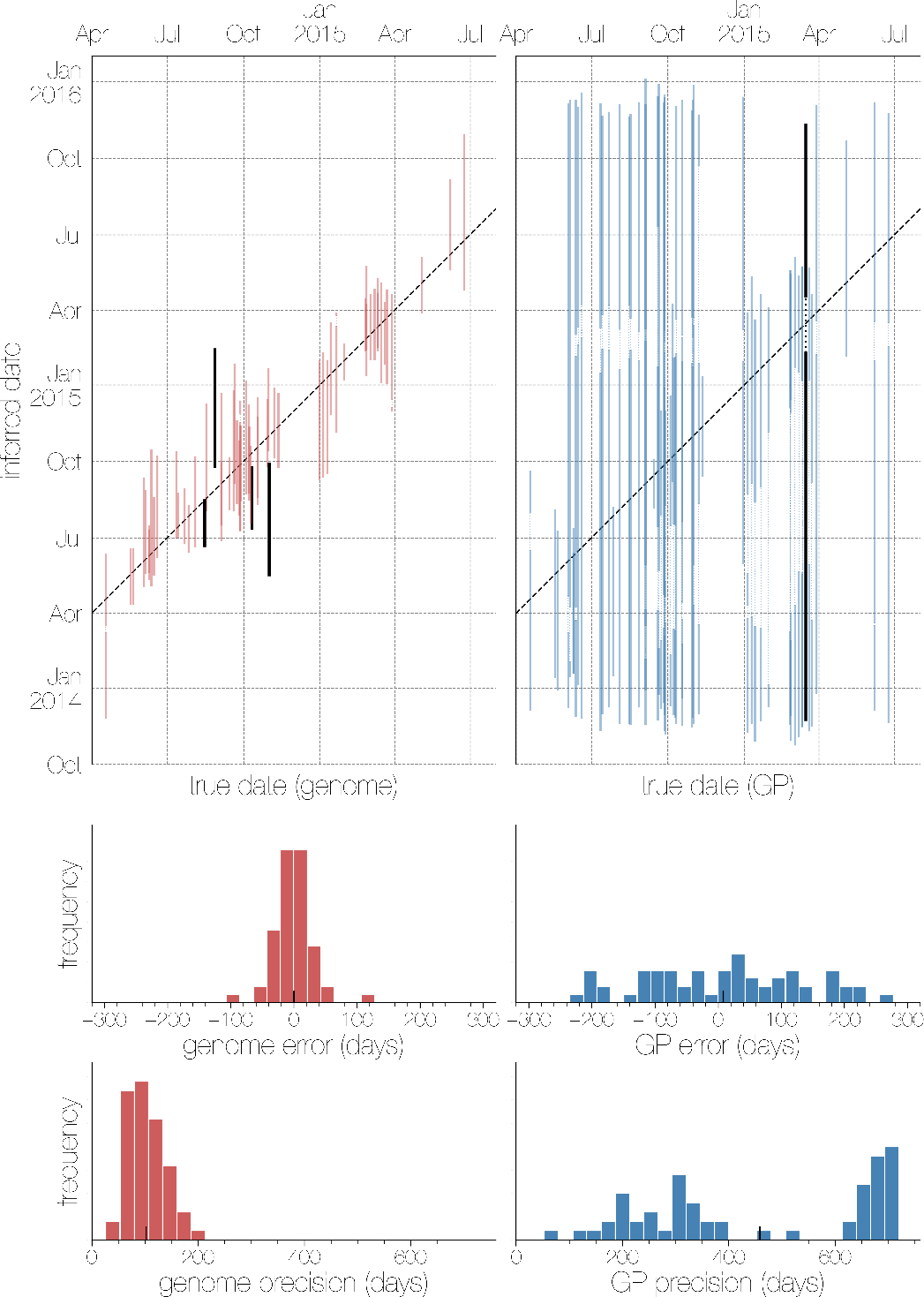
Masked tip date inference from genomes (left) and GP sequences (right). Inferred collection dates in the masked set based on genomes (red, left) and GP sequences (blue, right). Each vertical line corresponds to the 95% highest posterior density for the inferred tip date (y-axis), coloured red (genome) or blue (GP) if it falls within the true collection date (x-axis) and black otherwise. Dashed diagonal line indicates the 1-to-1 line. A histogram of residual errors (accuracy) between mean posterior date estimate and true date for each masked tip is shown in the second row. Third row shows the histogram of confidence interval widths for date estimates (precision).

### Migration model is strongly informed by tip dates and locations

Differences between genomic and GP datasets are clear and dramatic when looking at both phylogenies (Figure 1) and masked date inference (Figure 2), but less pronounced when trying to infer the location of a masked tip (Figure 3). Although locations are correctly inferred more often and with greater support in genomic sequences compared to just the GP gene, there are numerous tips whose locations are not correctly inferred even from genome sequences (Figure 3 and Figure S5 for maximum likelihood equivalent). This might reflect the nature of these parameters of interest, since phylogenies and date inference ultimately draw information from mutation accumulation via relatively straightforward models of sequence evolution with limited parameter space. In contrast, migration processes are far more complicated and nuanced without a *de facto* standard for modelling, though continuous time Markov chain (CTMC) approaches are widely used (Lemey et al., 2009) with most advanced methods relying on generalised linear models without excessive over-parameterisation (Faria et al., 2013; Lemey et al., 2014; Dudas et al., 2017). Despite the lack of strong contrast in power to infer masked locations between genomes and GP sequences, cross entropies indicate better performance with complete genomes (6054.631 nats) than with GP (9905.726 nats).

**Figure 3.**
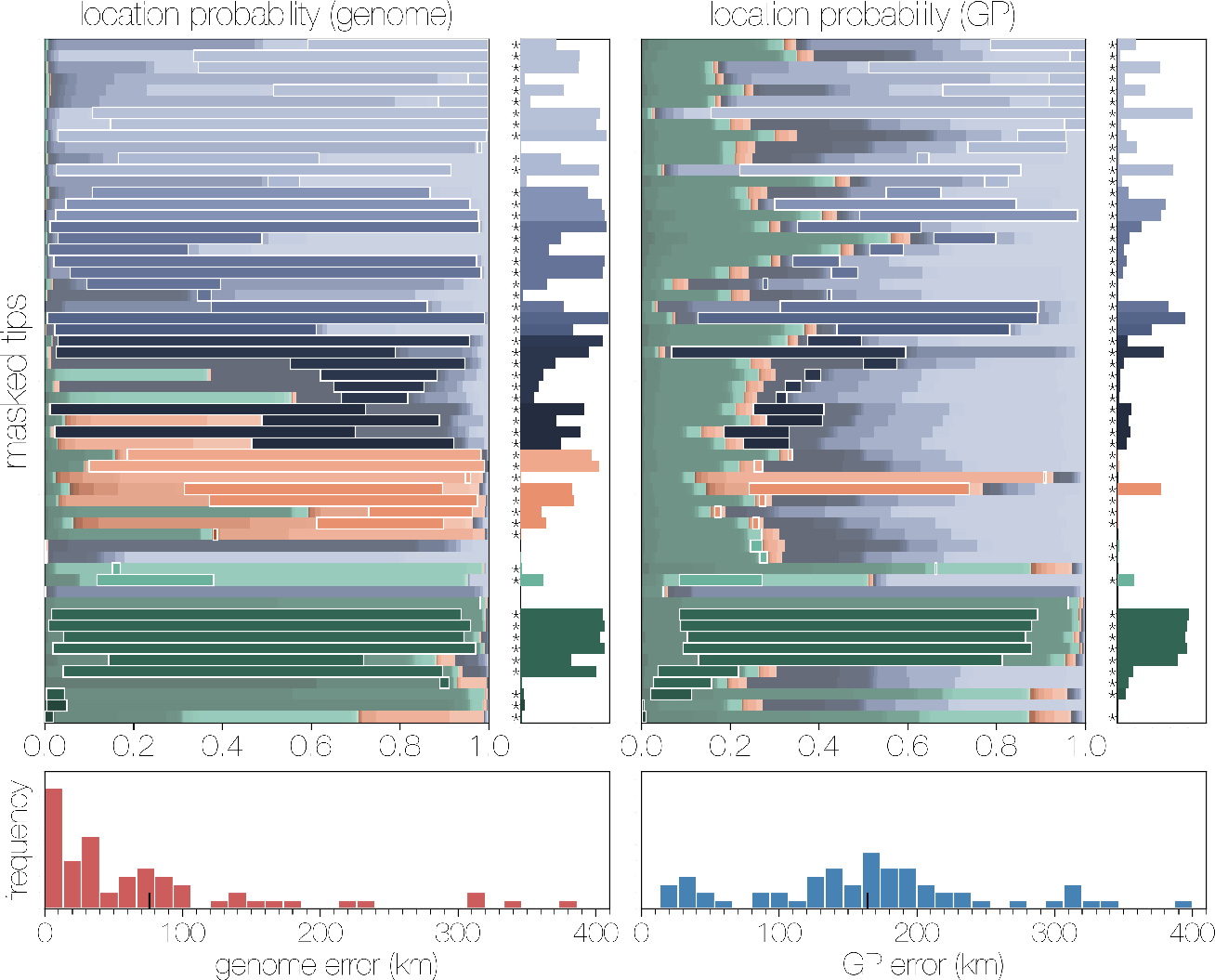
Masked tip location inference from genomes (left) and GP sequences (right) Horizontal bars indicate the posterior distribution of masked tip locations, coloured by country (Sierra Leone in blue, Liberia in red, Guinea in green) and location (lighter colours indicate administrative divisions lying towards west of the country). The correct location of each tip is outlined in white, with the smaller plot to the right showing only the posterior probability of the correct location. Bars marked with an asterisk indicate cases where the correct location is within the 95% credible set. Posterior distribution of probability-weighted distances between population centroids of inferred and correct locations is shown at the bottom with mean indicated by a tick mark.

Similarly, locations are inferred correctly more often with complete genomes than with GP sequences, where the maximum probability location (*i.e.* the model’s best guess) matches the truth, where using complete genomes results in 0.540 probability of guessing correctly compared to 0.286 probability for GP (for a calibration of both models see Figure S6). The model makes these guesses with more certainty too, where the mean probability of the true location is 0.482 with genomes and 0.219 with GP, and mean probability of best guess (*i.e.* maximum probability) is 0.680 and 0.396, respectively. We also calculated the great circle distances between the population centroids of true and each predicted location weighed by probability, which should ideally be 0.0 (0 km distance multiplied by probability of 1.0). The mean of these distances across masked tips are 75.886 kilometres for genomes compared to 164.309 km for GP sequences.

In addition to assessing how well tip locations can be inferred from genetic information we also looked at how well historical patterns were reconstructed from sequence data. To accomplish this we looked at the posterior distribution of ancestral locations of lineages that gave rise to four sequences in the data. The four lineages were chosen for their well-characterised histories in the broader epidemic as well as complexity of migration. One of these is strain ‘14859_EMLK’, a virus descended from the initial sweep of Ebola virus through Sierra Leone via Kailahun and Kenema districts which eventually ended up in Conakry (Guinea) from where it jumped back into Sierra Leone late in the epidemic where it was sequenced (Arias et al., 2016). ‘EM_004422’, which migrated through Kailahun district of Sierra Leone too, but then spread into Liberia from which it spilled back into Guinea via Macenta prefecture before ending up in Kissidougou (Guinea) where it was sequenced (Carroll et al., 2015). Similar to other lineages, ‘MK3462’ was part of the sweep through Sierra Leone, though unlike others remained in-country, migrating to the environs of Freetown and then Bombali district where it was sequenced (Arias et al., 2016). ‘PL5294’, unlike others, belonged to a lineage that was not part of the Sierra Leonean sweep and instead part of an unusual and under-sampled lineage endemic to western Guinea (“lineage A”, (Carroll et al., 2015)), from where it spilled into Sierra Leone’s Kambia district late in the epidemic (Arias et al., 2016).

The histories of these four tips are, for the most part, reconstructed from both GP sequences and genomes consistently (Figure 4), likely as a result of additional information brought in by specifying tip dates and their collection locations. In addition, genomic data tend to concentrate the probability mass towards a single location at any given time, in contrast to GP sequences where several locations can be considered with non-negligible probabilities at numerous time points (Figure 4 and Figure S7). What is even more apparent is that without the additional information available when using complete genomes MCMC explores a wider variety of low-probability migration paths, as indicated by maps on the right of each plot in Figure 4. In the case of ‘EM_004422’, for example, a series of migrations through distant Conakry (western Guinea) are reconstructed with relatively high confidence from GP sequences, compared to shorter distance migrations that run through neighbouring Liberia reconstructed from genomes.

**Figure 4.**
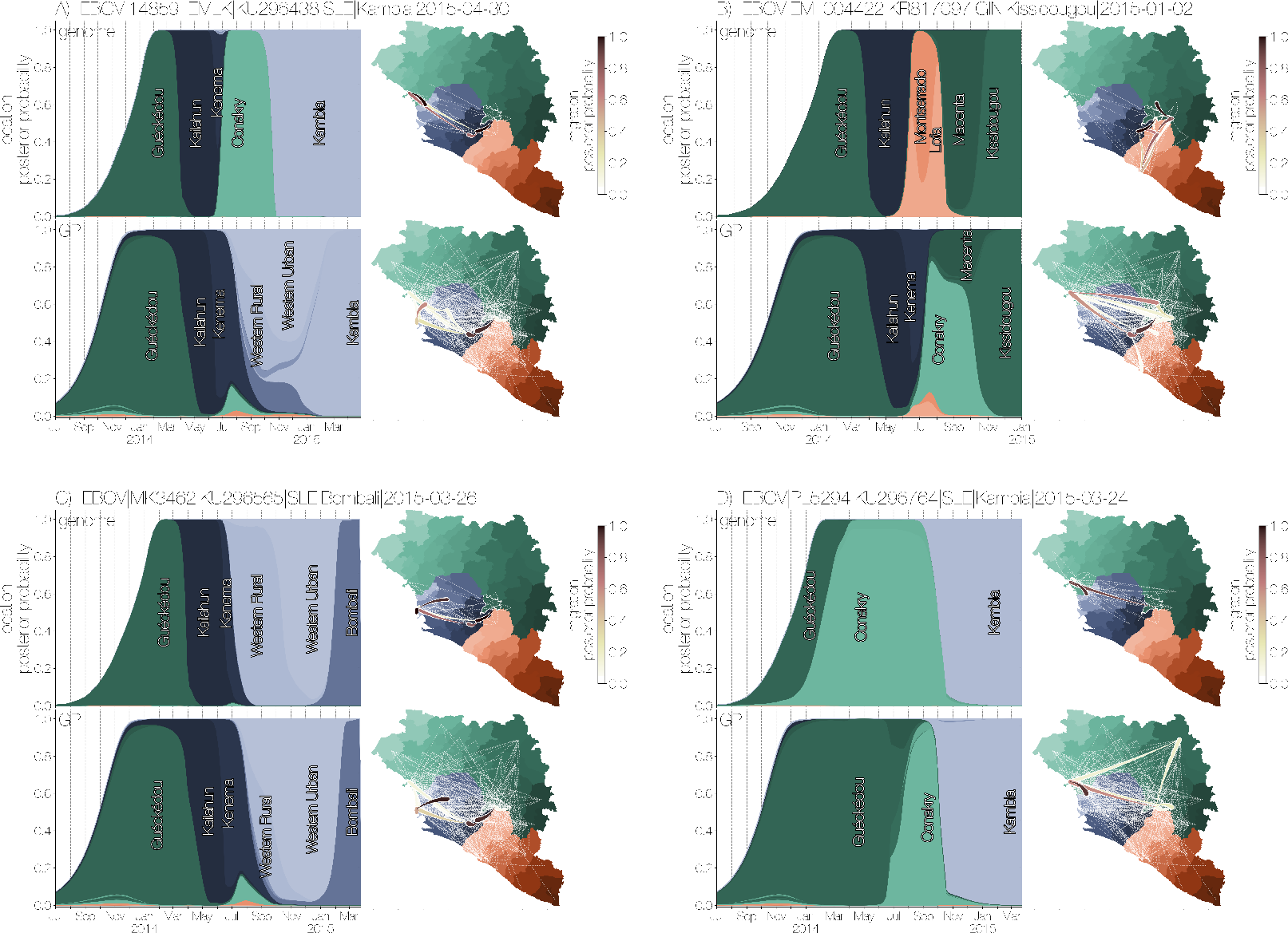
Posterior traces of ancestral locations and posterior migrations for four Ebola virus lineages from genomes (top) and GP sequences (bottom) The inferred ancestral branch location is logged at time points along the path from selected tips to the root of the tree, across the posterior distribution of trees. The smoothed trajectories are an indication of where and when a lineage that gave rise to a particular tip is inferred to have existed. Maps on the right show migration events that are inferred to have taken place, coloured by their posterior probability, with migrations with <0.05 posterior support are shown as dotted white lines. All lineages trace back to Guéckédou prefecture of Guinea (white outline in the map), where the original zoonotic transmission event occurred near the Guinean border with Sierra Leone and Liberia. Some lineages are also descended from an early spillover event into Sierra Leone.

Despite markedly reduced information content for both total number of sequences (>1600 to 600) and additional loss of information in GP (≈90% fewer sites) sequences, the same core correlates of migration are recovered for both datasets in the generalised linear model (Figure 5) compared to previous findings using all available sequence data (Dudas et al., 2017). These are: population sizes at origin and destination locations, within country migration effect, and great circle distances, which are identified as strong predictors of migration with high (>50 Bayes factor, BF), albeit not categorical, support (Figure 5). Four other migration predictors for the GP dataset have support >5 BF and <15 BF, which are international and national border sharing, Liberia-Guinea asymmetry, and index of temperature seasonality at origin. Of these Liberia-Guinea asymmetry and international border sharing are also found to be good predictors of migration in genomic data, though confusingly Liberia-Guinea asymmetry has the opposite correlation sign with GP sequence data. Apart from this deviation, predictors for both genome and GP gene datasets mostly have the same sign and very similar effect sizes. As mentioned previously (Figures 1 and 4), this suggests substantial amounts of information being derived from collection dates and locations of tips rather than genetic information. The reduction in total numbers of sequences, as well as reduced phylogenetic information in the GP dataset appears to enable the migration model to explore combinations of predictors that would otherwise be confidently excluded with complete genomes and thus a larger number of predictor matrices is included in the migration model with low probabilities. The differences between genomic and GP data, though seemingly small (*e.g.* Figure 3) is more pronounced when looking at total entropy of inclusion probabilities: 1.285 nats for genome data, and 2.688 for GP sequences.

**Figure 5.**
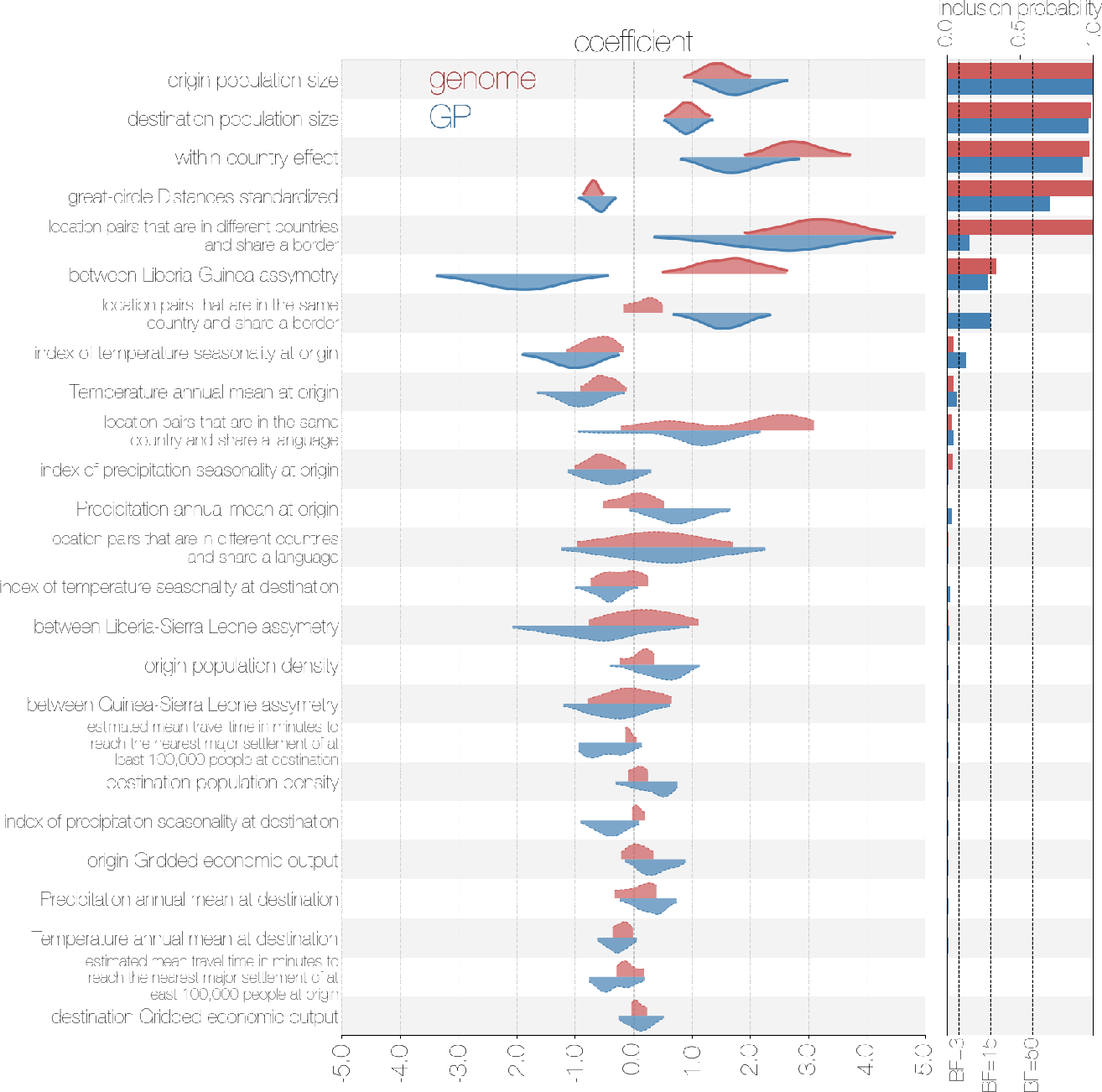
Correlates of migration identified from genomes (red) and GP sequences (blue). Effect size and direction of correlation between predictor matrices and migrations are shown as half violin plots, where the top kernel density estimates (in red) are derived from genomes and bottom kernel density estimates (in blue) are derived from GP sequences, conditioned on the predictor matrix being included in the model. Kernel density estimates of coefficients where predictors have <3 Bayes factor support are outlined in dashed lines. Posterior inclusion probabilities are shown on the right (red for genomes, blue for GP sequences) with appropriate Bayes factor cutoffs indicated by dashed lines.

### Temporal resolution

As discussed in the introduction, the mean waiting for a mutation is 1/*RL* (Equation 2) and depends on the rate at which mutations arise and are sampled by sequencing (evolutionary rate, *R*) and number of sites under observation (alignment length, *L*). Since 1/*RL* defines a linear relationship between rate *R* and length *L* mean waiting times for mutation can be reduced by an increase in either *R* or *L*. In order to double temporal resolution one can either double the evolutionary rate *R* or double the alignment length *L*. The former is generally outside the researchers’ control, though genomic regions evolving at a faster rate exist in many pathogens. How much faster smaller regions evolve will depend on forces of population genetics, such as ability to recombine with respect to the rest of the genome (strength of Hill-Robertson effect (Hill and Robertson, 1966)), as well as positive selection or functional constraints. It is thus unlikely that significantly higher rates will offset the reduction in resolution caused by focusing on a very small genomic region. Extending the region that is sequenced, on the other hand, is often trivial outside of resource-limited areas and can dramatically improve temporal resolution.

To help researchers intuit the impact of sequence length and evolutionary rate on temporal resolution we show the relationship between evolutionary rate and alignment length in determining mean waiting times until a mutation is observed in Figure 6. In addition to theoretical expectations we also show where a variety of viral pathogens fall along the two axes - estimated evolutionary rates with uncertainty intervals on the y-axis and alignment length on the x-axis. Sub-genomic alignments shown in Figure 6 include the small hydrophobic (SH) gene of mumps virus (Cui et al., 2017) and glycoprotein (GP) sequences of Ebola virus analysed in this study, as well as sequences of two human influenza A viruses - genome of subtype H1N1/09 (Hedge et al., 2013), and haemagglutinin sequences of subtypes H1N1/09 (Smith et al., 2009) and H3N2 (Rambaut et al., 2008). With respect to temporal resolution influenza A virus HAs are expected to acquire a mutation every one to two months on average, compared to around three to six months for Ebola virus GP and over a year for mumps virus SH. Though all of these sequences are from (-)ssRNA viruses, SH and GP genes are part of a single non-recombining RNA genome (Chare et al., 2003), whereas HA genes of influenza A viruses are encoded on their own segment which can be unlinked from their genomic background via reassortment. Because Ebola and mumps virus genomes do not recombine, their polymerases may have been selected for higher fidelity due to Hill-Robertson effect (Hill and Robertson, 1966).

Complete genomes, on the other hand, occupy parameter space that implies that a new mutation occurs on average every month or every few weeks. This is achieved through having more sites rather than substantial differences in evolutionary rates, which differ only marginally with respect to subgenomic fragments. Despite this, no virus is expected to acquire mutations faster than about once per week on average, and the two genomes with highest predicted temporal resolution - MERS-CoV and H1N1/09 - are difficult to analyse due to recombination and reassortment, respectively, though advances are being made in modelling reticulate evolution (Vaughan et al., 2017). The inverse relationship between observed evolutionary rate and sequence length is similar, but not the same as the relationship between virus genome sizes and mutation rates, where high mutation rates and large genome sizes lead to substantial deleterious mutation load (Pybus et al., 2007; Gago et al., 2009). This upper limit on mutation waiting times set by optimal evolutionary rates is what we refer to as the temporal horizon - population processes with inverse of rate (*i.e.* waiting time) less than the rate at which a pathogen acquires mutations will not be captured with high fidelity by currently existing methods. The exact relationship between mutation waiting times and rates of processes will, of course, be complicated by the presence of co-circulating lineages, site-wise rate heterogeneity, and choice of model for population processes of interest.

**Figure 6.**
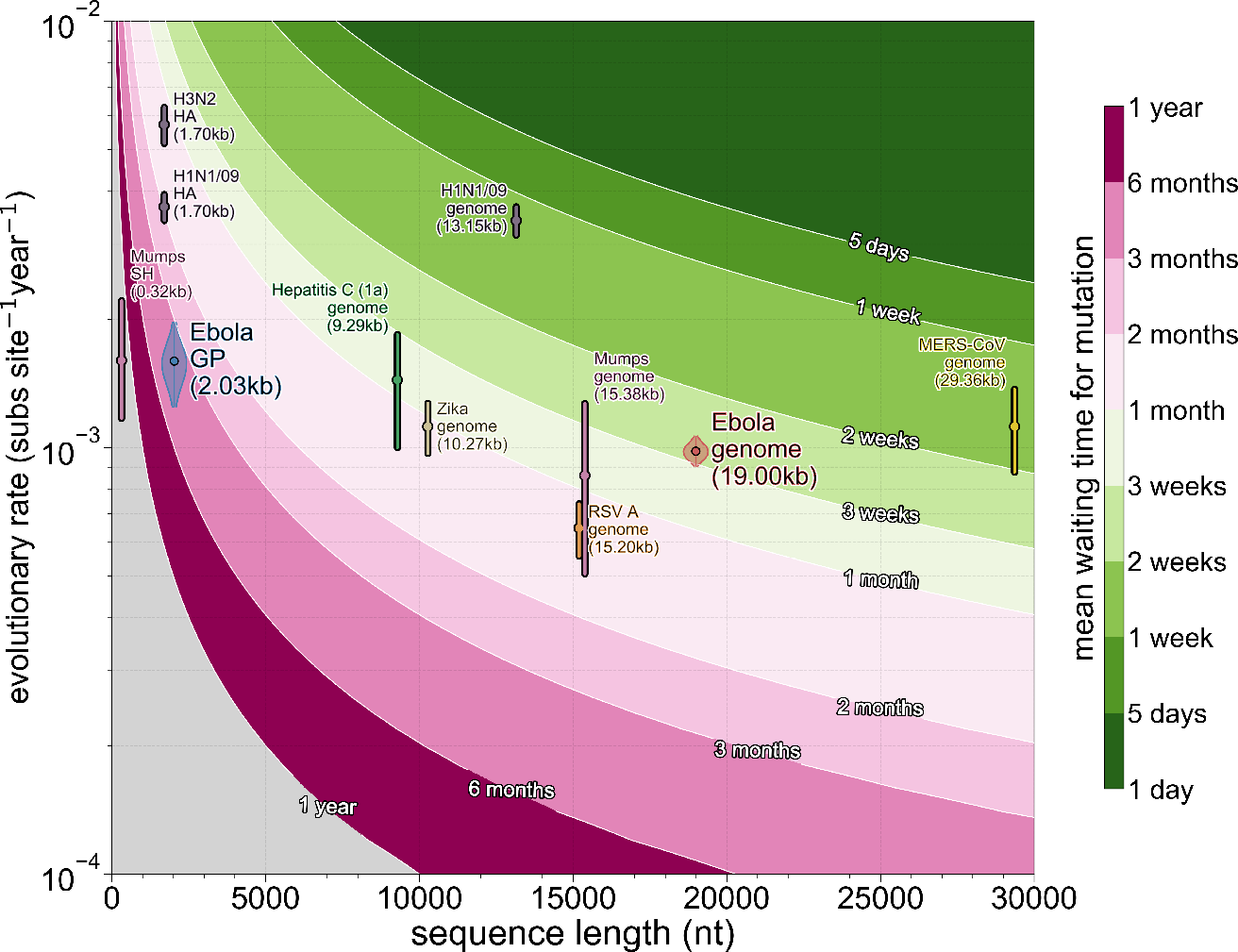
Mean waiting times for a mutation as a function of alignment length and evolutionary rate. Contours correspond to mean waiting times under a given combination of alignment length and evolutionary rate. Genomes and barcode genes for a variety of viruses are shown with reported evolutionary rate confidence intervals (vertical lines), including analyses of Ebola virus genomes (red violin) and GP sequences (blue violin) reported here. Most genomes occupy parameter space implying temporal resolution of a mutation once every month or so, with mumps virus as an exception. Sub-genomic fragments on the other hand are expected to have mean mutation waiting times of more than a month.

## Conclusions

### Theoretical considerations

For studies focused on temporal dynamics of pathogens over shorter periods of time the waiting time for a mutation should ideally be smaller than the inverse of the rate at which a process of interest occurs. Serial interval is often of most interest usually and has been addressed previously (Campbell et al., 2018; Grubaugh et al., 2019), but migration or cross-species transmission rates could also exceed the critical temporal resolution threshold if sequences are assigned to compartments that are too small. It is likely that this resolution limit will be improved greatly in the future by including additional information, either some aspects of a known transmission tree or, more likely, pathogen variation at the within-individual level, where variant sharing between two or more individuals is evidence of their linkage in a transmission cluster. Much like evolutionary rates, these methods might encounter biological limits outside of researchers’ control, however.

In addition to emphasising the need to sequence complete pathogen genomes, we also hope that our study imparts the interpretation of pathogen evolutionary rates as primarily a parameter indicating temporal resolution of sequence data, rather than a parameter of particular biological relevance. There have been previous incidents were a misunderstanding of the relationship between evolutionary rates and alignment length has been used to argue that low within-outbreak divergence in Ebola virus GP during the outbreak in Kikwit (Democratic Republic of Kongo) in 1995 was evidence of “genetic stability” (Rodriguez et al., 1999). What is far more likely to have taken place, however, is the phenomenon we show with our GP data (Figures 1 and S1), where even after more than two years of the West African epidemic the GP gene is too short to accumulate appreciable numbers of mutations. Higher reported evolutionary rates early in the West African epidemic (Gire et al., 2014) have also been misreported as having biological meaning, though not by the original study (Holmes et al., 2016; Rambaut et al., 2016), and arose through intense sequencing of a single transmission chain where mildly deleterious viral variants might not have been purged by purifying selection. We hope that our study clarifies that evolutionary rates are primarily a parameter of statistical resolution, rather than of evolutionary forces, and on their own are not sufficient to correctly interpret molecular clock data. Ideally, in the future sequence length and elapsed time will be included next to evolutionary rate estimates in order to transparently communicate statistical power available for analysis.

There is an additional Bayesian phylogenetic argument to be made in favour of using complete genomes. Molecular clock phylogenetics often relies on Markov chain Monte Carlo sampling to approximate the posterior distribution of phylogenetic trees (Yang and Rannala, 1997). Sequences which fall into polytomies in substitution phylogenies (*i.e.* well-defined common ancestry, but no indication of exact branching order) are particularly problematic, since plausible temporal phylogenies can be reconstructed in the absence of mutations. The branching order of such clades in time trees will be determined via tree and molecular clock rate priors, since no information about branching order can be recovered from the sequences themselves. There are over 34 million possible rooted trees for a set of 10 sequences, but many of these might not be visited during MCMC if, for example, sequences are collected over time and effective population size (*N_e_*) is low. Nonetheless, MCMC is particularly inefficient at sampling tree topologies for identical sequences (Whidden and Matsen, 2015), since increasing the number of identical sequences leads to expansion of search space without adding additional information that could constrain the search. Until reliable methods are developed and standardised the current solution is to reduce the numbers of identical sequences going into temporal MCMC analyses.

### Practical considerations

As well as temporal resolution concerns raised previously there are practical issues to consider when sequencing pathogens. Although many pathogens have established “barcode” genes or regions (Towner et al., 2004), some do not. This can easily lead to different groups sequencing different pathogen genes by chance or choice, as has happened with Ebola virus previously, where GP (Georges-Courbot et al., 1997), a short fragment of the polymerase (Leroy et al., 2005), or nucleoprotein (Rouquet et al., 2005) were sequenced, which is not necessarily a problem when sufficient complete genomes are available to bridge information between disparate regions and appropriate methods of analysis are used (Dudas and Rambaut, 2014). Sequencing complete pathogen genomes, in addition to providing the best possible resolution temporally in terms of mutation content (Figure 6), also ends up aiding in standardising data between studies, in the sense that a sequenced genome is a complete unit of data and there is nothing more to be done for sequence data except gathering better metadata.

It is also worth considering that the lifetime of sequence data extend beyond publication. Most scientific studies are designed with specific questions in mind that guide how data are collected and analysed to improve the researchers’ ability to detect differences. This makes combining data across studies with different goals (and correspondingly different data and approaches to analysing them) challenging. Sequence data, on the other hand, only become difficult to combine when sequences are too diverged to reliably align or are too numerous to infer phylogenies in reasonable time. Since divergence levels are generally low within outbreaks (with exceptions (Andersen et al., 2015)), sequence data are often trivial to combine. More than that, including sequence data from previous studies can reciprocally contextualise both older and newer sequences (*e.g.* Mena et al. (2016)). What remains problematic is determining and standardising additional data pertaining to the sequences themselves (“metadata”) in a way that makes sequence data easy to use by other groups. Whilst date and location of collection are widely reported and often of most interest, non-standard encodings of both are seen on public databases.

### Stating the obvious

It is not at all surprising that reducing the number of alignment columns by 10% from nearly 19,000 nucleotides that comprise the entire Ebola virus genome down to around 2,000 nt of the GP gene can result in loss of information, even if this shorter region evolves at a markedly faster rate. Here, we have quantified this loss of information via several methods: raw phylogenetic resolution (Fig 1), molecular clock signal (Fig 2), and aspects of migration model (Figs 3, 4, and 5).

In most cases biological aspects of the data, such as precise branching order and molecular clock resolution, suffer from severe loss in temporal resolution (Figure 2), whereas modelling of non-biological aspects of the data, *i.e.* migration, tend to be more robust (Figures 3 and 5). This is very likely to be caused by temporal and geographic, rather than genetic features of the sequence data (Boskova et al., 2018). A clustering of sequences from a particular location collected over a short period of time is likely to be a genuine outbreak cluster within a wider epidemic and in the absence of genetic information phylogeographic models tend to group sequences by location. This might explain why in many cases when comparing analysis results between genome and GP datasets statistical power in migration model remains disproportionately high, despite retaining only 10% of available sites and mutations and results between the entire >1600 genome data set (Dudas et al., 2017) is very similar to the reduced data set analysed here. On a similar note, case numbers alone have been used to recover a gravity-like model for the spread of Ebola virus in West Africa (Kramer et al., 2016) previously, further arguing that the clustering of cases in time and space contains sufficient information about the movement of Ebola virus in West Africa. The overall conclusion from our study, as well as others (Wohl et al., 2018), is that sequencing short genomic regions, instead of whole genomes is an ill-advised practice for investigating infectious disease outbreaks in any appreciable detail across relatively short timescales.

## Methods

### Sequence data

A publicly available dataset of 1610 Ebola virus genomes sequenced by various groups (Baize et al., 2014; Gire et al., 2014; Park et al., 2015; Carroll et al., 2015; Kugelman et al., 2015; Ladner et al., 2015; Simon-Loriere et al., 2015; Tong et al., 2015; Arias et al., 2016; Smits et al., 2015; Quick et al., 2015) and systematised in Dudas et al. (2017) was filtered to remove sequences where over 1% of the genome sequence was ambiguous or the precise location down to administrative division was not available, leaving 943 genomes. A set of 600 viral genomes were randomly sampled from the filtered dataset of 943 high quality genomes. Of the 600 genomes that were chosen for analysis 10% (60 genomes) were chosen for masking, where for all subsequent analyses both the date and location were considered as unknown and inferred as latent variables. Date inference was constrained to the period 2013 December 01 to 2015 December 01, corresponding roughly to the presumed beginning of the epidemic in late 2013 and its end in autumn of 2015. Another dataset was generated by extracting the glycoprotein GP coding sequence (with padding inserted into the polymerase slippage site to bring it in-frame) from the complete genomes dataset, resulting in an alignment 2031 nucleotides long.

### Bayesian analyses

Both GP and genome datasets were analysed in BEAST v1.8.10 (Suchard et al., 2018) under the generalised linear model (GLM) described previously (Faria et al., 2013; Lemey et al., 2014; Dudas et al., 2017) to infer the migration model. Sites in both GP and genome alignments were partitioned into codon positions 1, 2, and 3, with the genome analysis also including a partition comprised of non-coding intergenic regions. Each partition was assigned an independent HKY+Γ_4_ (Hasegawa et al., 1985; Yang, 1994) substitution model. A relaxed molecular clock (Drummond et al., 2006) with an uninformative prior on the mean (Ferreira and Suchard, 2008) of the log-normal distribution was used as the clock model. A flexible skygrid tree prior (Gill et al., 2013) was used to infer estimates of effective population size across 100 evenly spaced points in time starting 1.5 years prior to the collection of the most recent sequence to the date of the most recent sequence.

Both analyses (genome and GP) were set to run for 500 million states, sampling every 50,000 states and run three (genome) or seven (GP) times independently. Due to limited computational resources many analyses did not complete the full run and so for full genomes only 136.5, 8.62, and 143.8 million states were sampled, though after combining independent chains effective sample size (ESS) values are nearly the recommended 200. Similarly for GP only two MCMC analyses ran their allotted 500 million with others running to 259.9, 253.9, 255.8, 261.65, and 261.5 million states. Unlike complete genome MCMC analyses, GP analyses exhibit relatively poor ESS values even after combining seven independent chains, indicative of bad mixing in the absence of additional data contained in complete genome sequences. Convergence issues and appropriate burn-in values were assessed with Tracer v.1.7 (Rambaut et al., 2018), where 50 million states from every analysis (genome and GP) was discarded as burnin, with GP data additionally subsampled down to a quarter of the sampled states.

Posterior distributions of inferred tip dates for the masked set were logged during MCMC and 95% highest posterior density intervals were computed using a custom Python script, due to multi-peaked posterior distributions after combining independent analyses. Posterior distributions of trees were summarised as maximum clade credibility (MCC) trees using TreeAnnotator (Suchard et al., 2018). Inferred posterior probabilities of masked tip locations were recovered from MCC trees. Ancestral location probabilities were recovered via a modified version of baltic’s samogitia.py (https://github.com/evogytis/baltic/blob/master/samogitia.py) across 200 equally spaced time points between mid-2013 and beginning of 2016.

### Maximum likelihood analyses

RAxML (Stamatakis, 2014) was used to infer maximum likelihood phylogenies for genome and GP datasets under the same partitioning as described for Bayesian analyses: three codon position partitions for GP and genome, with genomes having an additional partition for intergenic regions under independent GTR+CAT substitution models. Trees were rooted in TreeTime according to best r^2^ value for root-to-tip against collection date regression with the 2 year constraint used for masked tips described earlier. A temporal phylogeny with marginal reconstruction of most likely dates for masked tips was carried out in TreeTime (Sagulenko et al., 2018) as well. Ancestral sequences at internal nodes of the clock-rooted RAxML topology were inferred using TreeTime under an HKY model (Hasegawa et al., 1985) of evolution. Ancestral location states were inferred in TreeTime using a continuous time Markov chain model identical to the one used by Lemey et al. (2009) without the Bayesian stochastic search variable selection. We also repeated many of the analyses under a maximum likelihood model in TreeTime (Sagulenko et al., 2018), like inference of masked tip dates (figure S4) and locations (figure S5).

### Error computation

For figure 2 mean absolute errors were computed as

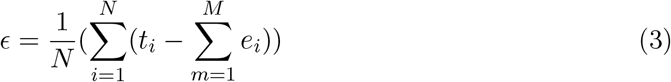

Where N is the number of masked tips, *t*_*i*_ is the true date of the *i* th masked tip, *e*_*i*_ is the estimated date of the *i*th masked tip, and M is the number of states sampled from the posterior distribution.

For figure 3 errors expressed in units of distance were calculated as

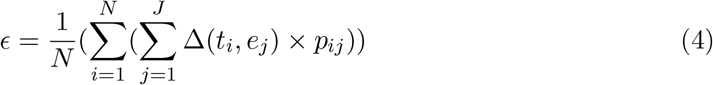

Where N is the number of masked tips, J is the number of locations in the migration model, Δ is great circle distance in kilometres, *t*_*i*_ is the coordinate of the population centroid of the true location of the *i* th masked tip, *e*_*j*_ is the coordinate of the population centroid of *j* th location, and *p*_*j*_ is the probability that the *i* th tip is in *j* th location.

Entropies for predictors shown in figure 5 and location probabilities in figure S7 were calculated as

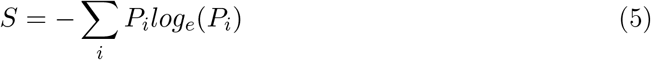

where *P*_*i*_ is the mean posterior inclusion probability of *i* th predictor matrix in the model for figure 5, and probability of *i* th location for figure S7.

Cross entropies for figure 3 were calculated as

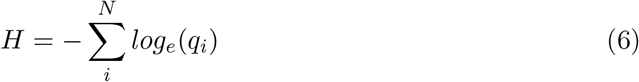

where N is the number of masked tips, *q*_*i*_ is the probability of the true location of the *i* th masked tip, which is assigned a probability of 0.0001 if the true location does not appear in the set of inferred possible locations (*i.e.* has probability 0.0) to avoid domain error.

### Data availability

Sequence data and all analytical code is publicly available at https://github.com/blab/genomic-horizon.

## Acknowledgements

We would like to thank Andrew Rambaut for useful discussion and advice. GD is supported by the Mahan postdoctoral fellowship from the Fred Hutchinson Cancer Research Center. TB is a Pew Biomedical Scholar and is supported by NIH R35 GM119774-01.

---

**Figure S1.**
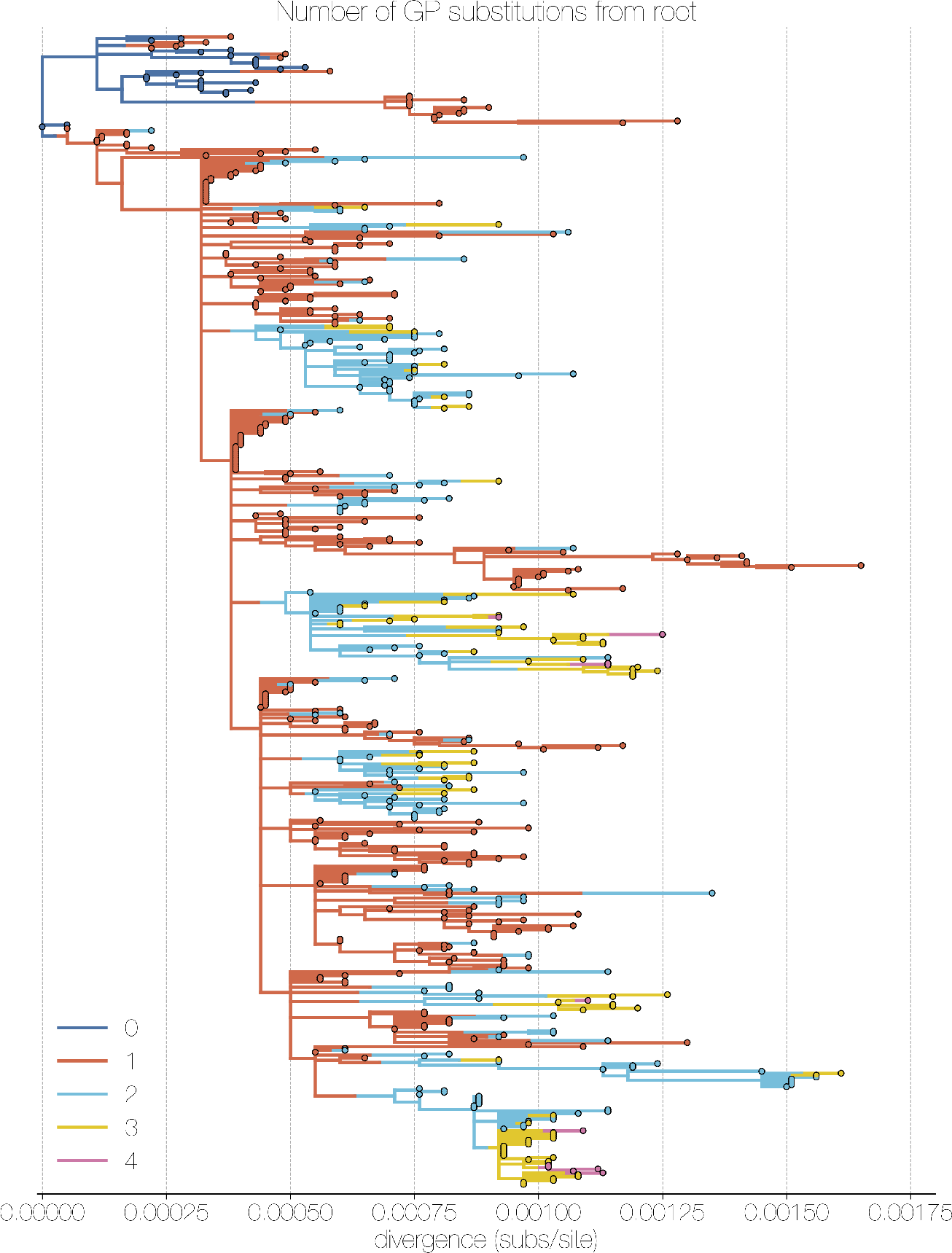
Whole genome maximum likelihood tree coloured by mutations occurring in GP. Colours indicate the cumulative number of mutations from the root occurring in the GP gene. Much of the clade resolution is lost when only considering mutations occurring in the GP gene, particularly in the already highly polytomic Sierra Leonean part of the phylogeny in red.

**Figure S2.**
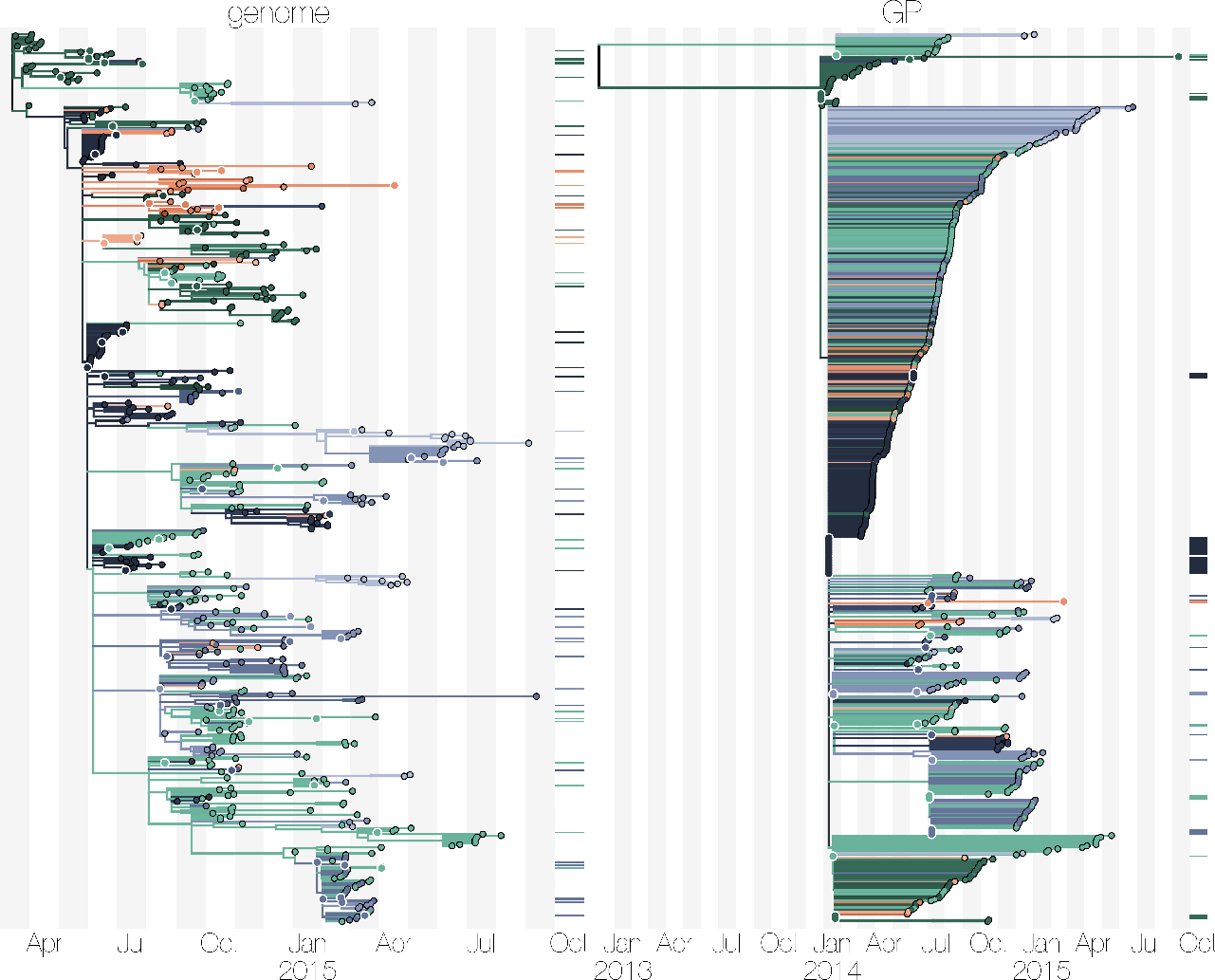
Maximum likelihood phylogenies of complete Ebola virus genomes (left) and GP sequences (right) with maximum likelihood ancestral location reconstruction. Trees were inferred in RAxML (Stamatakis, 2014) with ancestral state reconstruction performed in TreeTime (Sagulenko et al., 2018). Inferred phylogeographic patterns are for the most part consistent with Bayesian results presented in Figure 1 with severe loss of statistical power when using GP instead of genome sequences.

**Figure S3.**
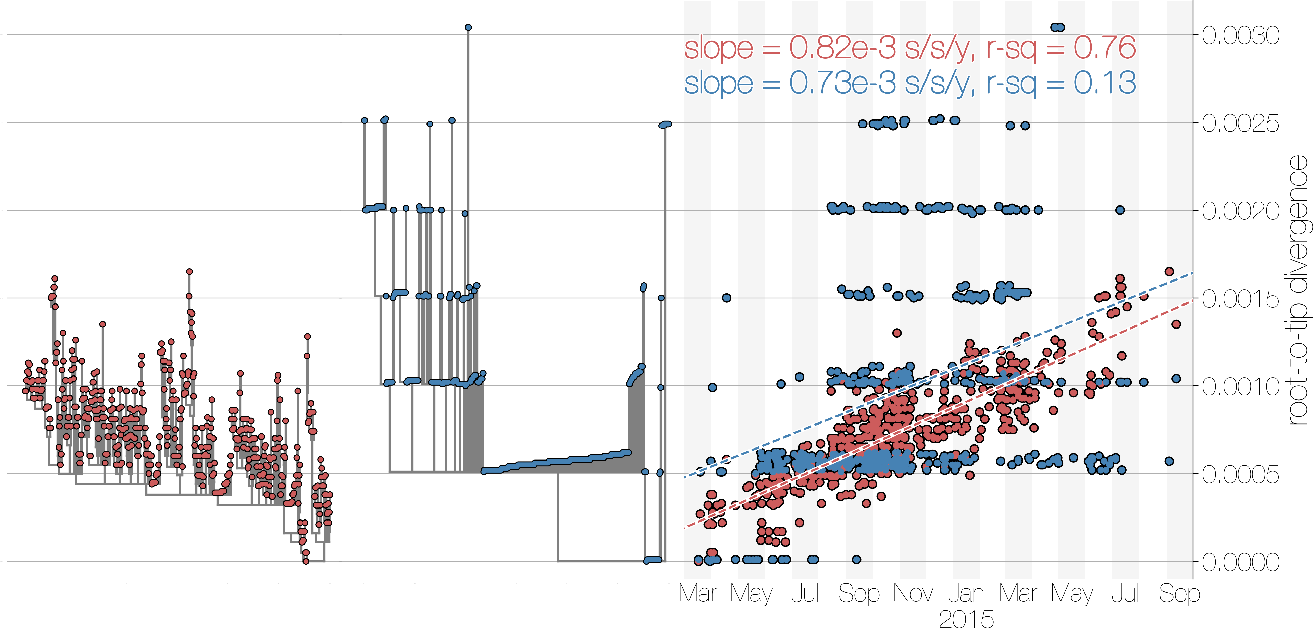
Root to tip regression for maximum likelihood trees of genome (red) and GP (blue) sequences. Linear regression of sequence collection dates against distance from the root gives evolutionary rate estimates (slope of the regression) at 0.82×10^−3^ and 0.73×10^−3^ substitutions per site per year, respectively. Despite similar rates the correlation between collection dates and divergence from root is far better using genomes (*r*^2^ = 0.76) than GP sequences (*r*^2^ = 0.13).

**Figure S4.**
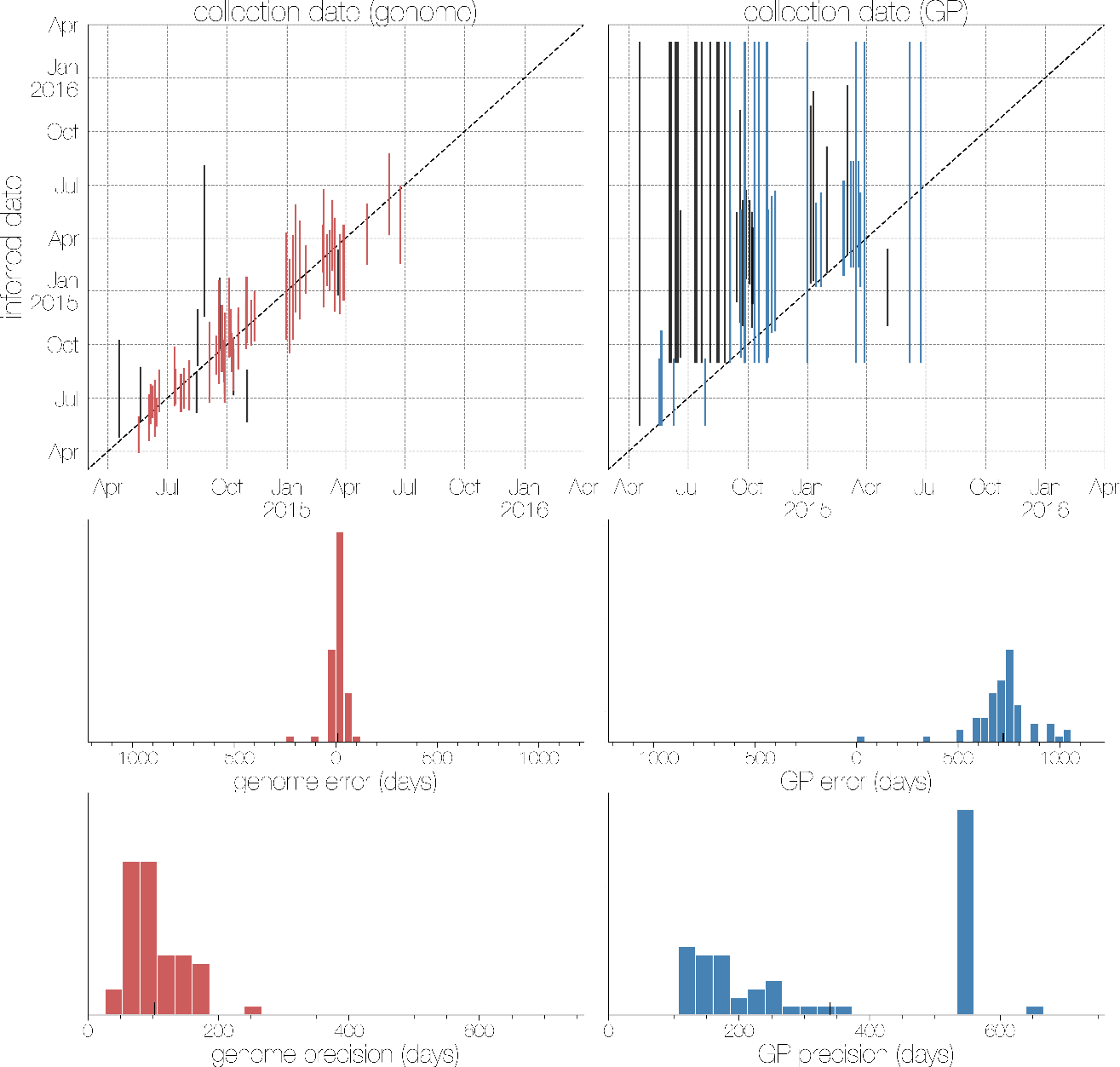
Maximum likelihood inference of masked tip dates from genomes (red, left) and GP sequences (blue, right) using TreeTime. Vertical bars indicate the 95% confidence interval for marginal reconstruction of masked tip dates plotted against their true dates. Tip dates where the 95% confidence interval excludes the true value are shown in black.

**Figure S5.**
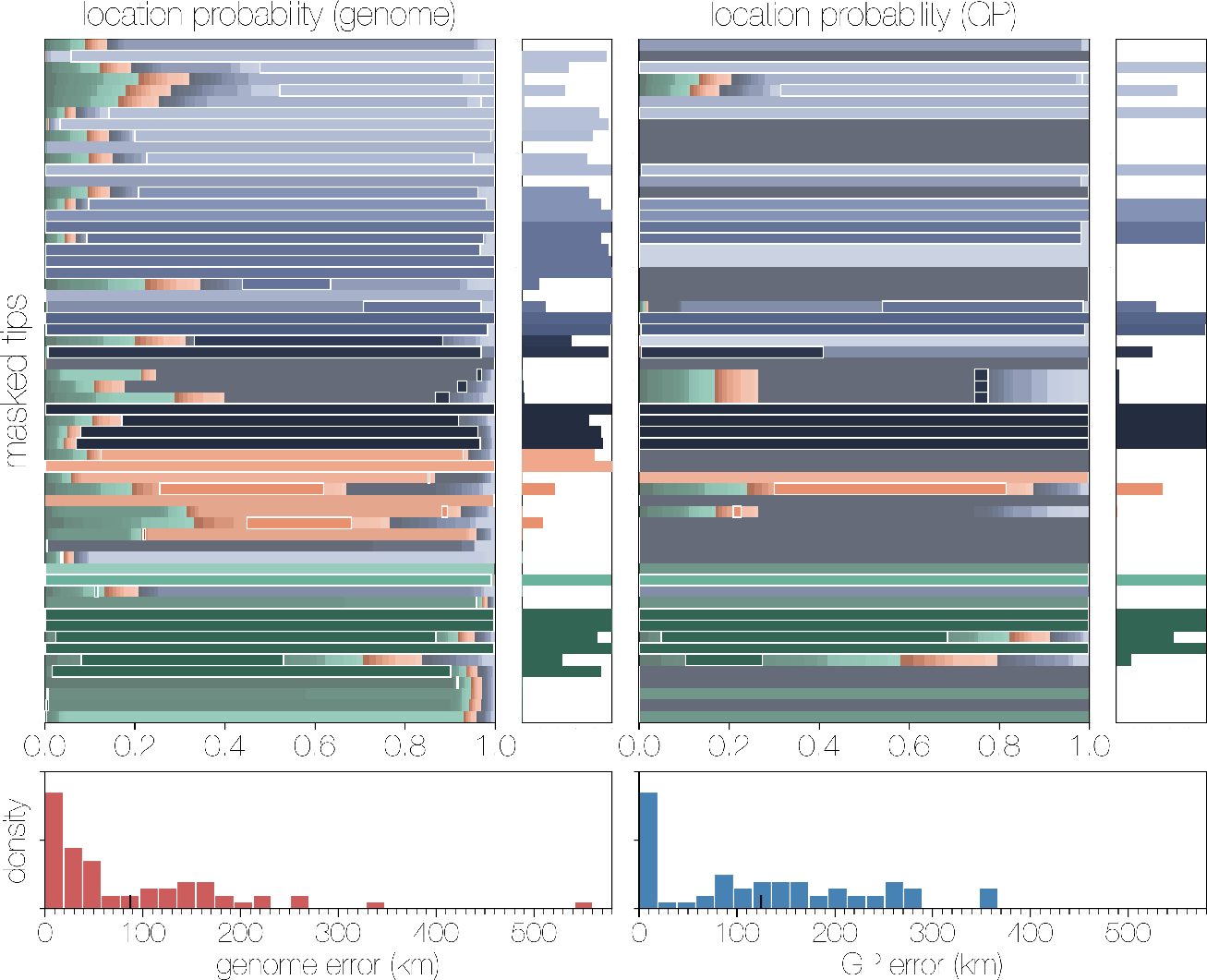
Maximum likelihood inference of masked sequence location from genomes (left) and GP sequences (right) via a CTMC model implemented in TreeTime. Unlike the Bayesian GLM where probabilities are spread across different locations, the CTMC gives categorical support to one location. Genomes still perform better in terms of correct guess (0.432 probability that best guess location is true location for genomes versus 0.259 for GP), cross entropy (12012.800 nats for genome versus 24397.109 nats for GP) and mean probability-weighted great circle distance between true location population centroid and estimated location population centroid (87.568 km for genome versus 124.909 km for GP).

**Figure S6.**
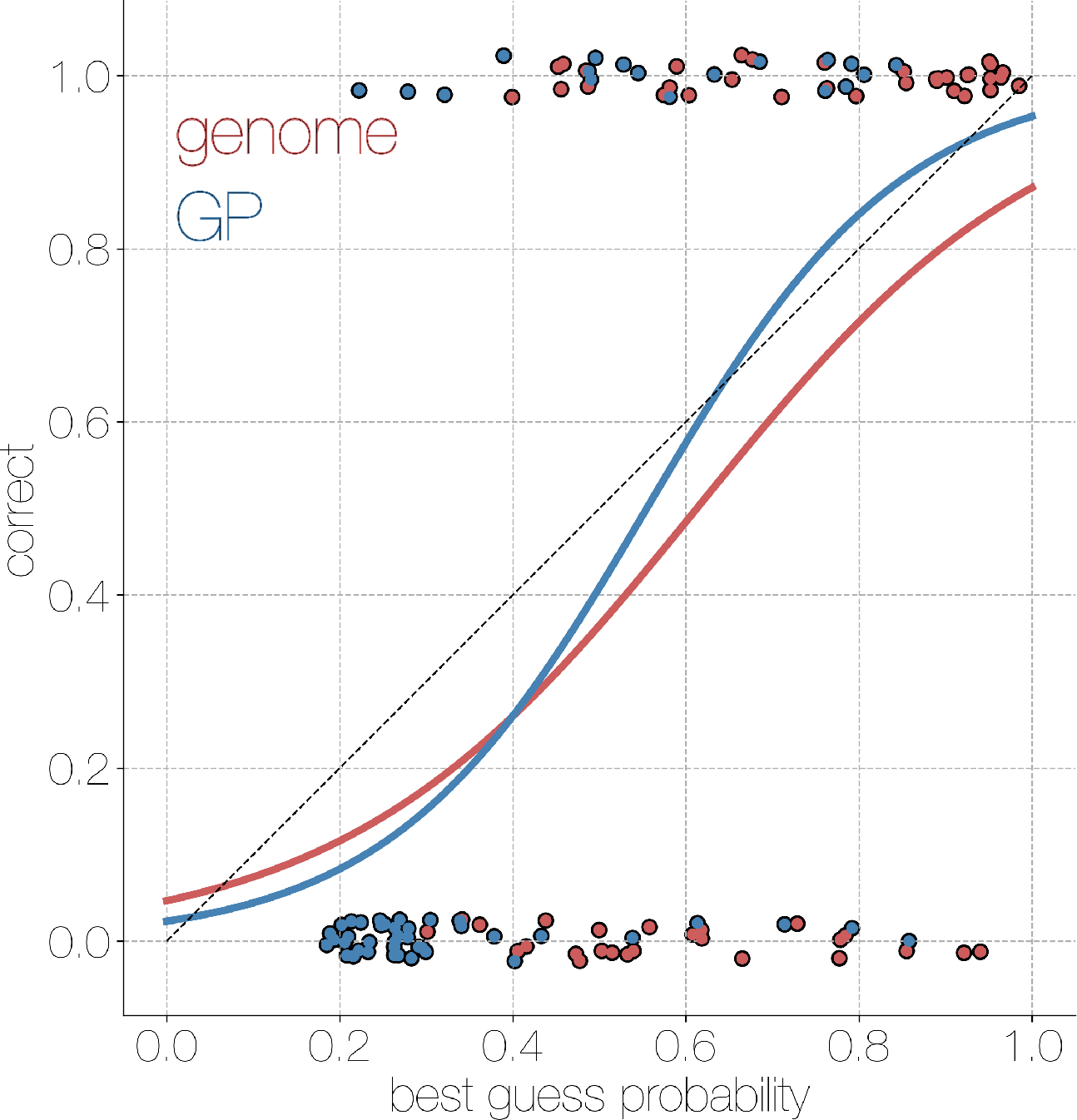
Calibration curve for phylogeographic model informed with genome (red) and GP (blue) sequences. Logistic regression of probability of the most likely location against whether it is correct or not for genome (red) and GP (blue) sequences with jitter introduced along the y axis to make points discernible. Overall performance of the phylogeographic model is comparable between genome and GP sequences, as indicated by sigmoid curves matching the 1-to-1 dotted line.

**Figure S7.**
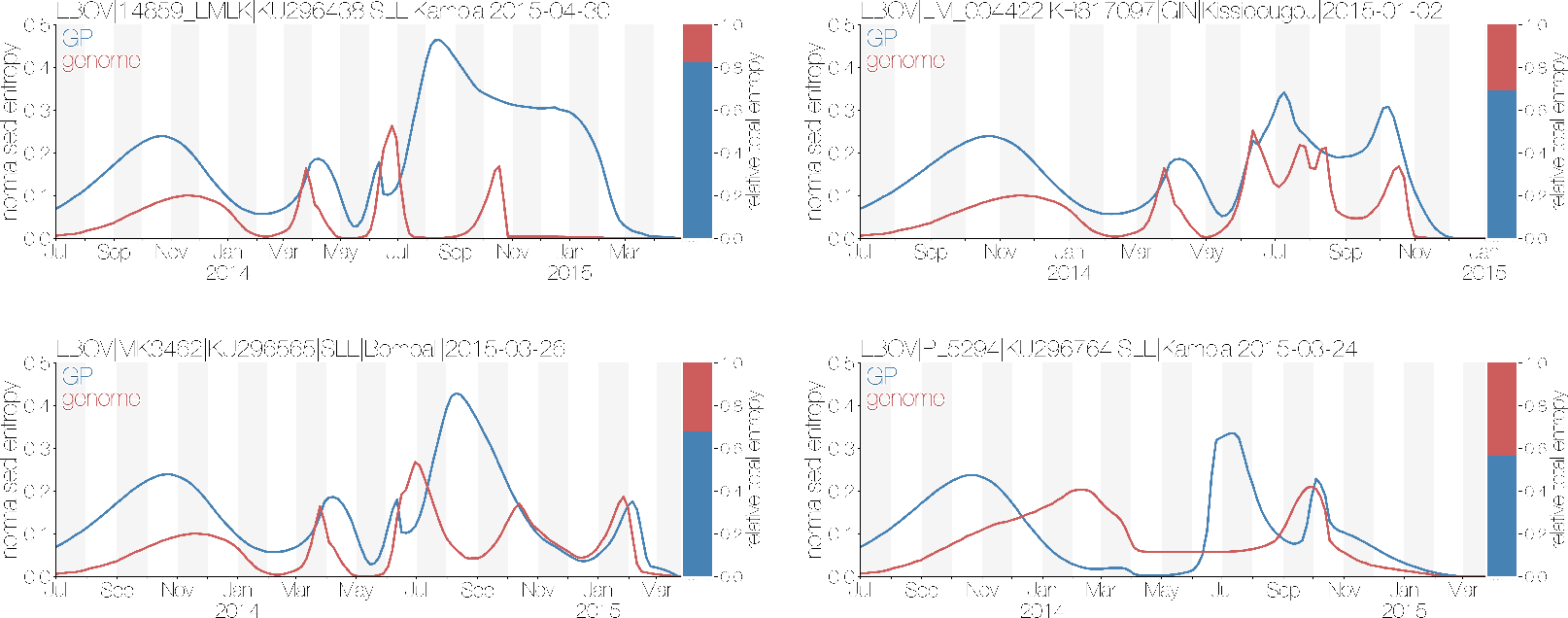
Entropies of posterior ancestral location reconstruction from genomes (red) and GP sequences (blue) for four tips. Ancestral state reconstructions from genomes typically have lower entropies relative to reconstructions derived from GP sequences, indicating better certainty in location assignment at any given time. Red and blue bars at the end of the plot indicate relative cumulative entropies of genome and GP sequence reconstructions, respectively.

